# Adipocyte MKK3 Increases in Human Obesity protecting against Insulin Resistance by p38β activation

**DOI:** 10.1101/2025.05.19.654793

**Authors:** Edgar Bernardo, Nuria Matesanz, Jorge-Luis Torres, Leticia Herrera-Melle, Luis Leiva-Vega, Alfonso Mora, Maria Elena Rodriguez, Lourdes Hernández-Cosido, Rubén Nogueiras, J. Simon C. Arthur, Angel R Nebreda, Roger J Davis, Miguel Marcos, Guadalupe Sabio

## Abstract

Obesity is a major global health concern, and a key predisposing factor for insulin resistance and type 2 diabetes mellitus. Adipocytes play a critical role in the development of obesity-induced insulin resistance, with several signaling pathways influencing insulin sensitivity. Among these, p38 MAP kinases are essential for adipose tissue physiology and the regulation of processes such as differentiation, thermogenesis, and inflammation. p38 activation is mediated by the upstream kinases MKK3 and MKK6 in response to inflammatory signals. While MKK6 inhibition promotes browning and thermogenesis and protects against obesity, the role of MKK3 remains unclear.

Here, we investigated the function of MKK3 in adipose tissue. In human adipose tissue samples, *MKK3* expression was positively correlated with body mass index (BMI) and negatively correlated with glycated hemoglobin, a marker of hyperglycemia.

Using whole-body and adipose-specific *Mkk3* and *p38β* knockout mice, we found that *Mkk3* activation in adipose tissue during obesity enhances insulin sensitivity. Mechanistically, adipose tissue from Mkk3- or p38β-deficient mice exhibited elevated basal p70S6K activity compared with wild-type controls. This increased p70S6K activity was linked to higher serine phosphorylation of insulin receptor substrate 1 (IRS1) and impaired insulin-stimulated Akt phosphorylation, contributing to worsened insulin resistance.

Collectively, our data suggest that activation of the MKK3/p38β signaling axis in adipocytes may protect against high-fat diet–induced insulin resistance and diabetes.

## Introduction

Obesity is a major risk factor for insulin resistance and type 2 diabetes mellitus. It is characterized by chronic, low-grade inflammation in adipose tissue (1), with increased immune cell infiltration and elevated production of inflammatory cytokines such as tumor necrosis factor α (TNFα) and interleukin 6 (IL-6) (2). Both cytokines have been shown to induce insulin resistance in adipocytes (3, 4) and systemically in distant organs (5). Although the mechanisms linking obesity and insulin resistance are not fully understood, the activation of stress kinases (SAPKs), including JNK and p38, has been implicated. This activation contributes to both local and systemic insulin resistance, particularly via inflammation in white adipose tissue (WAT) (5–7).

Indeed, the p38 pathway is activated in adipose tissue, where it regulates inflammation, adaptation, and remodeling (6). It is also activated in other tissues in patients with obesity, including T cells infiltrating adipose tissue (8). Distinct roles have been described among the four p38 MAPK isoforms: p38α, p38β, p38γ, and p38δ (9). p38α has been reported to mediate insulin resistance in TNFα-treated skeletal muscle and endothelial cells (10, 11). In myeloid cells, p38α controls TNF and IL6 production by macrophages, contributing to obesity-induced liver steatosis (12). Additionally, p38γ and p38δ influence muscle fiber composition and voluntary activity (13, 14), and promote neutrophil infiltration and steatosis in the liver during obesity(15–17).

Furthermore, p38α and p38β regulate IL-6 production in adipocytes (13), whereas p38α inhibits thermogenesis in brown adipose tissue (BAT) in contrast to p38δ, which promotes thermogenesis (8). The primary upstream kinases responsible for activating the p38 pathway in adipocytes are MKK3 and MKK6. Interestingly, deletion of both kinases in the myeloid compartment modulates the liver macrophage–hepatocyte–adipocyte axis, ultimately regulating fat thermogenesis (18).

Although it was thought that both MKK3 and MKK6 were able to activate all the p38 family members, recent studies have suggested that MKK3 and MKK6 do not activate all p38 family members equally, and their roles may not be redundant (19–21). Instead, MKK3 appears to selectively activate p38γ and p38δ, while MKK6 preferentially activates p38α and p38β (19, 22). This differential activation implies that MKK3 and MKK6 may play unique roles in adipocyte function and remodeling in obesity. For example, MKK6 has been shown to be involved in processes such as browning of white adipose tissue and thermogenesis (21), whereas MKK3 may regulate other aspects of adipocyte biology related to inflammation and insulin resistance through p38γ and p38δ.This specificity of MKK3 and MKK6 in activating different p38 isoforms suggests that they could have opposing or complementary effects on adipose tissue biology. Understanding this selective activation helps elucidate the distinct roles played by these MAPKs in metabolic diseases. Obesity is associated with increased MKK6 expression in human and mouse WAT (21). However, the regulation of MKK3 remains unknown. Using a cohort of human adipose tissue samples, we found that MKK3 expression directly correlated with BMI and blood leptin levels. In contrast, its levels are inversely correlated with the hyperglycemic marker glycated hemoglobin (HbA1c). To understand the physiological role of adipocyte MKK3 in regulating whole-body metabolism and insulin sensitivity, we generated mice with adipocyte-specific *Mkk3* deletion. We found that these mice developed insulin resistance in adipose tissue *in vivo* and in *Mkk3* KO-derived primary adipocytes. Evaluation of the p38 family members involved in this phenotype indicated that the phenotype was due, at least in part, to a decrease in p38β activation, as mice specifically lacking this kinase also developed insulin resistance. In both animal models, insulin resistance correlated with increased activation of p70S6K and serine phosphorylation of insulin receptor substrate 1 (IRS1). Hyperactivation of p70S6K in adipose tissue has been shown to induce insulin resistance and alter whole-body metabolism (23). Our data indicate that inhibition of the increased basal activation of p70S6K was sufficient to restore insulin sensitivity *in vitro*.

## Results

### *Mkk3* mRNA expression levels in white adipose tissue positively correlate with BMI and leptin serum levels in humans

Stress kinases in adipose tissue have been linked to both positive and negative effects of obesity (24). In fact, while lack of p38α in adipose tissue protects against obesity, depletion of p38δ results in increased brown adipose tissue generation, ameliorating obesity (9), suggesting distinct roles for the different components of the p38 pathway. MKK3 and MKK6 are the main activators of the p38 family members. Although deletion of MKK6 protects against obesity (21), the role of MKK3 remains unknown.

To evaluate the potential role of MKK3 in obesity-mediated insulin resistance in humans, we analyzed circulating levels of insulin, C-peptide, adiponectin, and leptin in the plasma of lean and obese subjects. As previously described, obese individuals presented with hyperinsulinemia, elevated circulating levels of C-peptide and leptin, and lower circulating levels of adiponectin than lean individuals (25, 26) (Fig. 1A-D). To investigate the role of adipose MKK3 in obesity-induced diabetes, we measured *MKK3* expression in visceral fat from a cohort of 71 subjects. Interestingly, *MKK3* mRNA levels positively correlated with BMI (Fig. 1E). Additionally, a positive correlation between adipose *MKK3* RNA and circulating leptin levels was observed (Fig. 1F), whereas no correlation was found with other metabolic parameters (Table 1). Conversely, *MKK3* expression was negatively correlated with HbA1c levels (Fig. 1G), suggesting that higher *MKK3* expression may contribute to maintaining lower glycemic levels in obese subjects. Finally, we found that adipose *MKK3* expression positively correlated with serum C-peptide levels, suggesting that patients with higher MKK3 expression also exhibit hyperinsulinemia (Fig. 1H).

**Figure 1.**
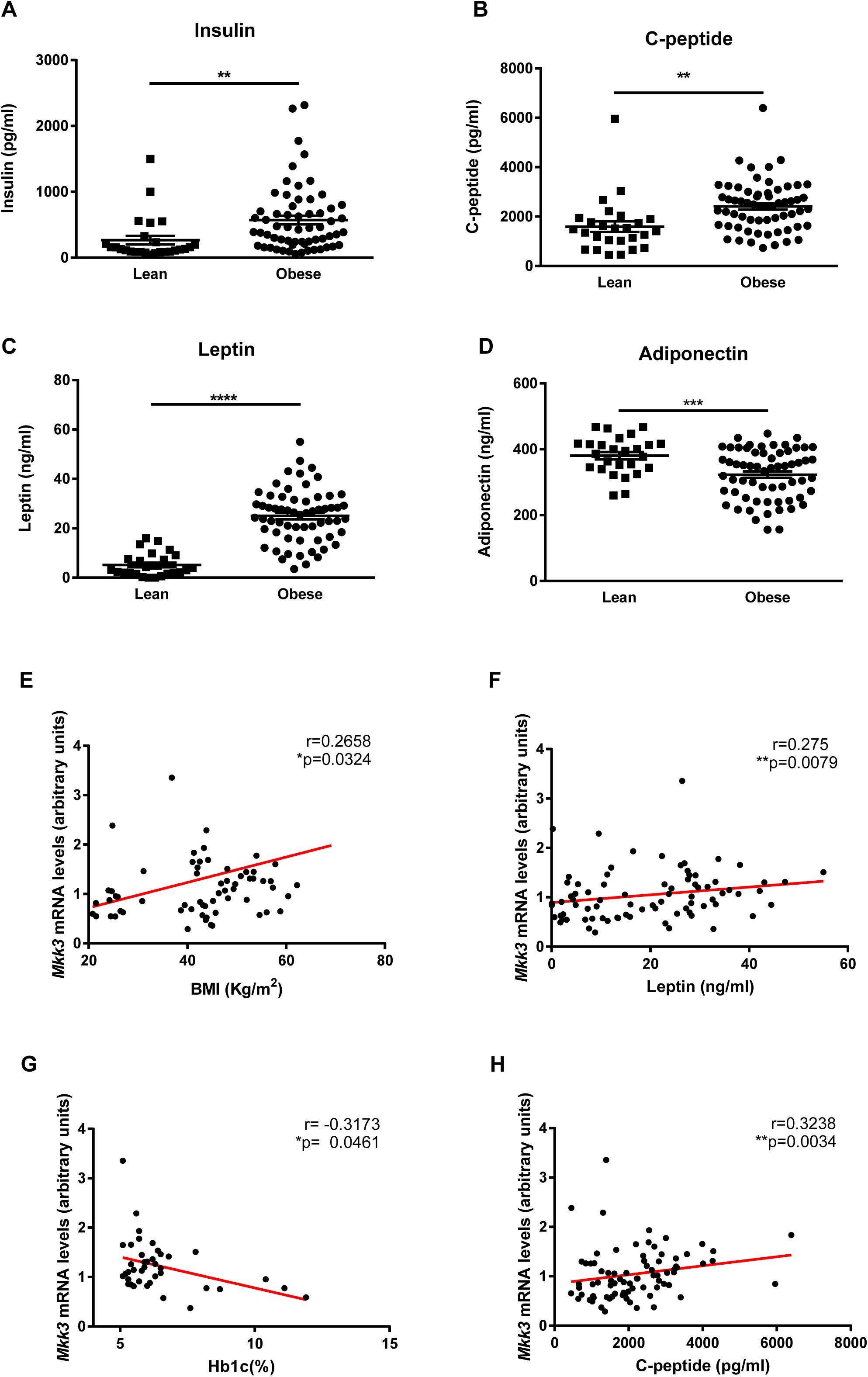
**A-D**. Increased levels of insulin (A), C-peptide (B), leptin (C), and reduced levels of adiponectin (D) in serum from lean and obese human individuals as measured by the Luminex test. Data are presented as mean ± SEM, n=18-53, Student’s *t-*test coupled with Welch’s correction, **p<0.01, ***p<0.001, ****p<0.0001. **E-H.** *Mkk3* expression in visceral adipose tissue from lean and obese human subjects positively correlated with BMI (E) and circulating levels of leptin (F) and C-peptide (G), and negatively correlated with Hb1Ac (H). Data are presented as discrete value comparisons between individuals and regression slope calculations, n=48-71, Spearmańs Rho, *p<0.05, **p<0.01.

### *Mkk3* deficient mice develop adipose tissue-dependent insulin resistance after HFD

Given that MKK3 upregulation in human adipose tissue might represent a compensatory response to obesity-induced diabetes, we tested whether MKK3 deficiency affects insulin signaling in *Mkk3* knockout mice fed a high-fat diet (HFD) for 12 weeks.

No differences in body weight, fasting or fed blood glucose levels, or glucose tolerance were observed between the genotypes during HFD feeding (Fig. 2A–D). However, whole-body insulin sensitivity testing revealed that after insulin injection, *Mkk3^-/-^* mice had significantly higher blood glucose levels than controls (Fig. 2E). This indicates that mutant mice exhibited a poorer response to insulin stimulation, suggesting that during obesity, insulin resistance is increased in peripheral tissues.

**Fig. 2.**
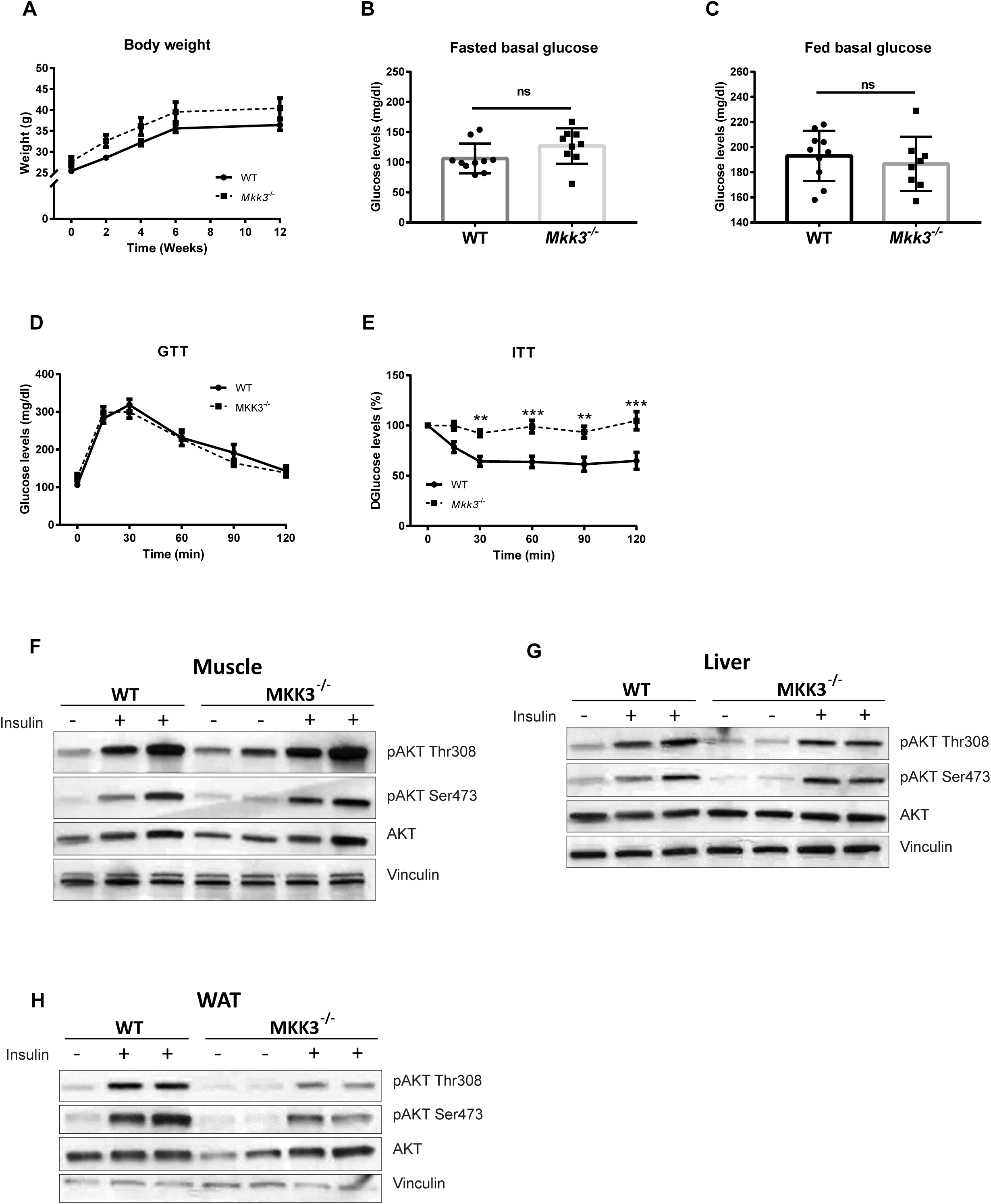
**A**. Body weight of 8 weeks old WT and *Mkk3^-/-^* mice fed an HFD for 12 weeks. Data are presented as the mean ± SEM, n=10, Two-way ANOVA coupled with Bonferronís post-test. **B, C.** Overnight-fasted (B) and fed (C) glycemic levels of WT and *Mkk3^-/-^* mice fed an HFD for 12 weeks showed no significant differences. Data are presented as mean ± SEM (n = 10), Student’s *t*-test coupled with Bonferronís post-test. **D.** Glucose tolerance test (GTT) of HFD-fed WT and *Mkk3^-/-^* mice. Data are presented as the mean ± SEM, n=10, Two-way ANOVA coupled with Bonferronís post-test. **E.** Insulin tolerance test (ITT) showing that *Mkk3^-/-^*mice are insulin intolerant after intraperitoneal injection of insulin. Data are presented as mean ± SEM of decrease with respect to initial glycemia calculated as a percentage, n=10, Two-way ANOVA coupled to Bonferronís post-test, **p<0.01, ***p<0.001. **F-H.** Immunoblots of AKT activation in muscle (F), liver (G), and white adipose tissue (WAT) (H). Fasted mice were treated with or without 1.5 IU/kg intraperitoneally injected insulin for 15 min.

Interestingly, despite their reduced insulin sensitivity, *Mkk3^-/-^*mice exhibited a significant reduction in fat mass as assessed by both MRI and direct tissue weighing (Fig. S1A–C). MRI analysis also showed increased lean mass, increased muscle mass, and unchanged liver mass (Fig. S1D–F). These findings indicate that Mkk3 may regulate not only insulin signaling but also body composition. The paradoxical combination of leanness and insulin resistance observed in *Mkk3^-/-^*mice suggests a strong insulin-resistant phenotype, despite reduced adiposity.

Insulin signaling assessment in different tissues showed no differences in the muscle and liver (Fig. 2F, G). However, insulin-stimulated AKT phosphorylation was reduced in white adipose tissue (WAT) of *Mkk3^-/-^* mice (Fig. 2H), indicating that adipose tissue-specific insulin signaling was impaired in HFD-fed *Mkk3^-/-^*mice.

### HFD-fed *Mkk3* deficient mice present increased leukocyte infiltration in white adipose tissue

Macrophage infiltration contributes to obesity-induced insulin resistance (27, 28). Histological evaluation showed increased infiltration of leukocytes surrounding the adipocytes, forming crown-like structures (CLS) in WAT from HFD-fed *Mkk3^-/-^* mice compared to control mice (Fig. S1G). Additionally, slight increases in adiposity were noted in the BAT and livers of *Mkk3^-/-^* mice. Immunofluorescence confirmed increased macrophage (F4/80+) and neutrophil (Ly6G+) infiltration in WAT (Fig. S1H). We also observed reduced perilipin staining in WAT from HFD-fed *Mkk3^-/-^* mice, a marker associated with CLS, and increased macrophage and neutrophil infiltration in WAT from HFD-fed *Mkk3^-/-^* mice, which was confirmed by immunofluorescence staining of F4/80 (red) and Ly6G (gray), respectively (Fig. S1H). Additionally, we observed a loss of perilipin (green), which was associated with the presence of CLS (27). These data suggest that either the stromal vascular fraction (SVF) or adipocytes might be responsible for the increased insulin resistance and decreased fat mass observed in the WAT of HFD-fed *Mkk3^-/-^* mice. Despite increased leukocyte infiltration, the mRNA levels of pro-inflammatory cytokines (including chemoattractants, Il-6, and TNF-α) in WAT were unchanged in HFD-fed *Mkk3^-/-^*mice (Fig. S1I–N).

### Insulin resistance in *Mkk3* deficient mice is independent of myeloid cells

To determine whether insulin resistance in *Mkk3^-/-^* mice was mediated by a lack of Mkk3 in myeloid cells, we generated bone marrow (BM) chimeras by transplanting BM from WT or *Mkk3^-/-^* donors into irradiated WT recipients, referred to as WT^WT^ or WT*^Mkk3-/-^*mice, respectively (Fig. S2A), and fed them an HFD for 12 weeks. No differences in weight gain, tissue weight, glucose tolerance, or insulin sensitivity were observed (Fig. S2B-H). These results indicate that *Mkk3* deficiency in the BM compartment did not affect HFD-induced insulin resistance.

To corroborate these results, conditional mouse models lacking *Mkk3,* specifically in the myeloid compartment *Lyz2-Cre+Mkk3^f/-^* (*Mkk3*^Lyz2-KO^) and *Lyz2-Cre+Mkk3^+/-^* (Lyz2-Cre) as control mice, were generated and fed HFD. No differences between genotypes were found in body weight, liver, muscle, WAT, BAT weight, or insulin sensitivity (Fig. S2I-N). All these data indicate that deficiency of *Mkk3* in the myeloid compartment might not play a role in obesity-induced insulin resistance, suggesting that this phenotype might be originated in an adipocyte autonomous specific manner.

### Adipocyte-specific *Mkk3* deletion induces insulin resistance and impaired insulin signaling in epididymal white adipose tissue

As HFD-fed *Mkk3^-/-^* mice exhibited exacerbated whole-body and adipose tissue insulin resistance, which is unlikely to be due to the deficiency of *Mkk3* in the myeloid compartment, we evaluated whether these effects would be adipose tissue-specific. To corroborate this hypothesis, mice lacking *Mkk3*, specifically in the adipose tissue, were generated. *Mkk3* deletion was observed in different tissues, indicating that MKK3 was depleted in the WAT (Fig. 3A).

**Fig. 3.**
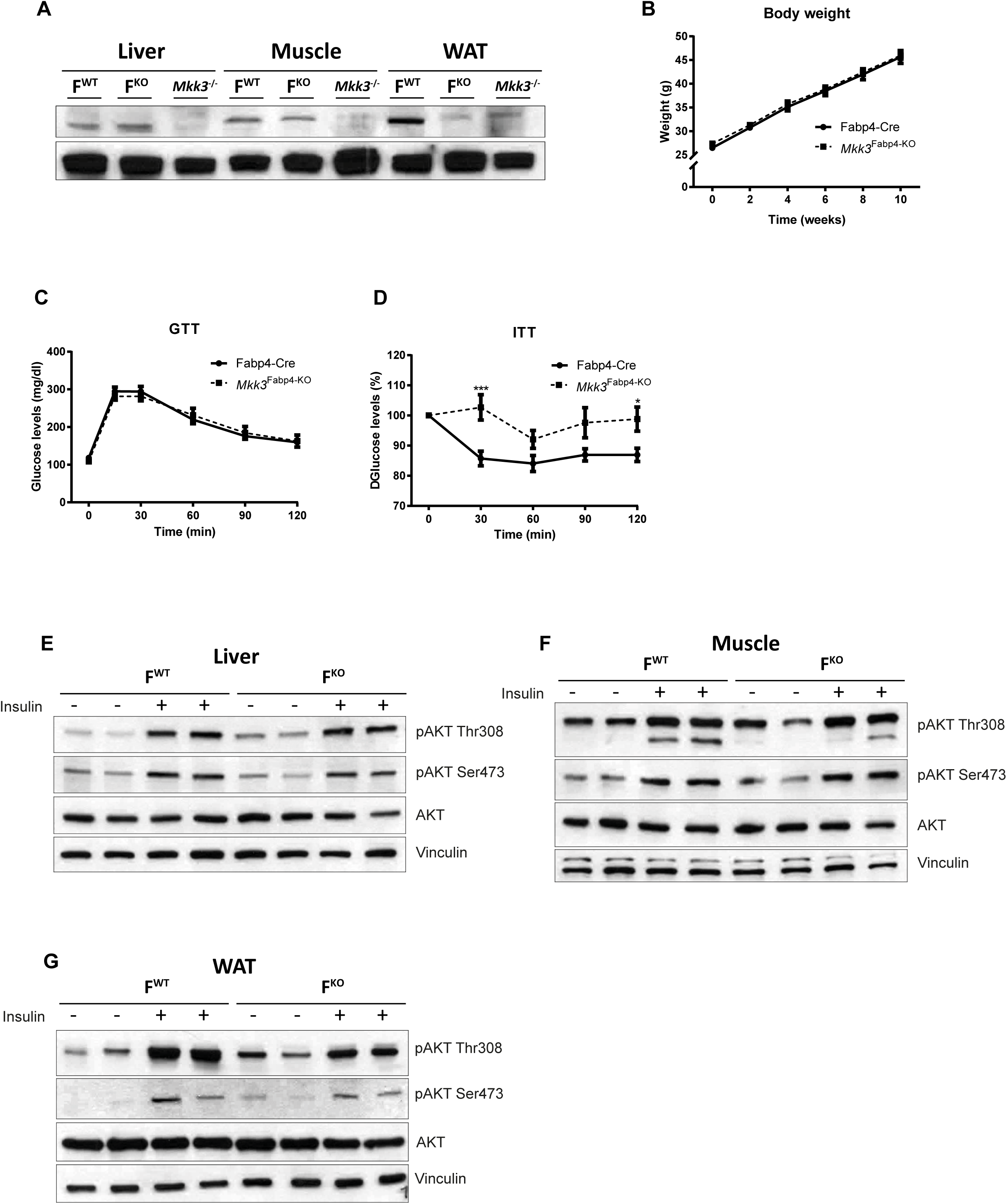
**A**. Western blot showing white adipose tissue-specific depletion of MKK3 in control (F^WT^) *Mkk3*^Fabp4-KO^ (F^KO^) mice **B.** Body weight of 12 weeks old Fabp4-Cre and *Mkk3*^Fabp4-KO^ mice fed an HFD for 10 weeks. Data are presented as mean ± SEM, n=8-11, Two-way ANOVA coupled with Bonferronís post-test. **C.** GTT in HFD-fed Fabp4-Cre and *Mkk3*^Fabp4-KO^ mice. **D.** ITT of HFD-fed Fabp4-Cre and *Mkk3*^Fabp4-KO^ mice showing increased insulin resistance of mutant mice. C and D, Data are presented as mean ± SEM, n=8-11, Two-way ANOVA coupled with Bonferronís post-test. *p<0.05, ***p<0.001. ITT data are presented as mean ± SEM of the decrease with respect to initial glycemia, calculated as a percentage. **E-G.** Western blot showing AKT activation specifically in liver (E), muscle (F), and white adipose tissue (WAT) (G) of HFD-fed *Mkk3*^Fabp4-KO^ mice. Fasted mice were treated with or without 1.5 IU/kg intraperitoneally injected insulin for 15 min.

*Mkk3*^Fabp4-KO^ and Fabp4-Cre mice, as controls, were fed an HFD, showing no differences in weight gain, tissue weight, or glucose tolerance between genotypes (Fig. 3B, C and Fig. S3A-D). However, when insulin sensitivity was assessed, HFD-fed *Mkk3*^Fabp4-KO^ mice showed impaired blood glucose clearance after insulin injection compared with control mice (Fig. 3D), indicating that adipose tissue-specific *Mkk3* deficient mice were more insulin resistant. We then analysed which tissue was resistant to insulin in these mice and observed that, similar to Mkk3^-/-^ mice, insulin-induced Akt phosphorylation in the liver and muscle was similar in both genotypes (Fig. 3E, F). WAT from *Mkk3*^Fabp4-KO^ mice showed reduced insulin-induced Akt phosphorylation at both Thr308 and Ser473 residues, indicating that WAT from *Mkk3*^Fabp4-KO^ mice presented higher insulin resistance than control mice (Fig. 3G).

The decrease in Akt phosphorylation in adipose tissue from *Mkk3*^Fabp4-KO^ mice, but not in the muscle and liver, and the resemblance with the same phenotype observed in HFD-fed germinal *Mkk3* deficient mice strengthened the hypothesis that the lack of *Mkk3* in adipocytes is sufficient to induce insulin resistance both in WAT and systemically, suggesting that *Mkk3* in adipose tissue plays a critical role in maintaining whole-body glucose homeostasis.

### Insulin resistance in WAT from *Mkk3^-/-^* mice is cell autonomous

Our previous results suggested that the lack of *Mkk3* in adipocytes might impair insulin signaling, leading to severe insulin resistance in HFD-fed animals. Thus, to elucidate the molecular mechanisms underlying this phenotype, differentiated mature adipocytes from WT and *Mkk3*^-/-^ mice were used (Fig. 4A). Oil Red O staining confirmed no differences between the WT and *Mkk3^-/-^* adipocytes in terms of differentiation and lipid accumulation (Fig. 4B). Quantification of *Adiponectin* mRNA expression levels also confirmed that adipocytes were fully differentiated, showing no differences in expression between genotypes (Fig. 4C).

**Fig. 4.**
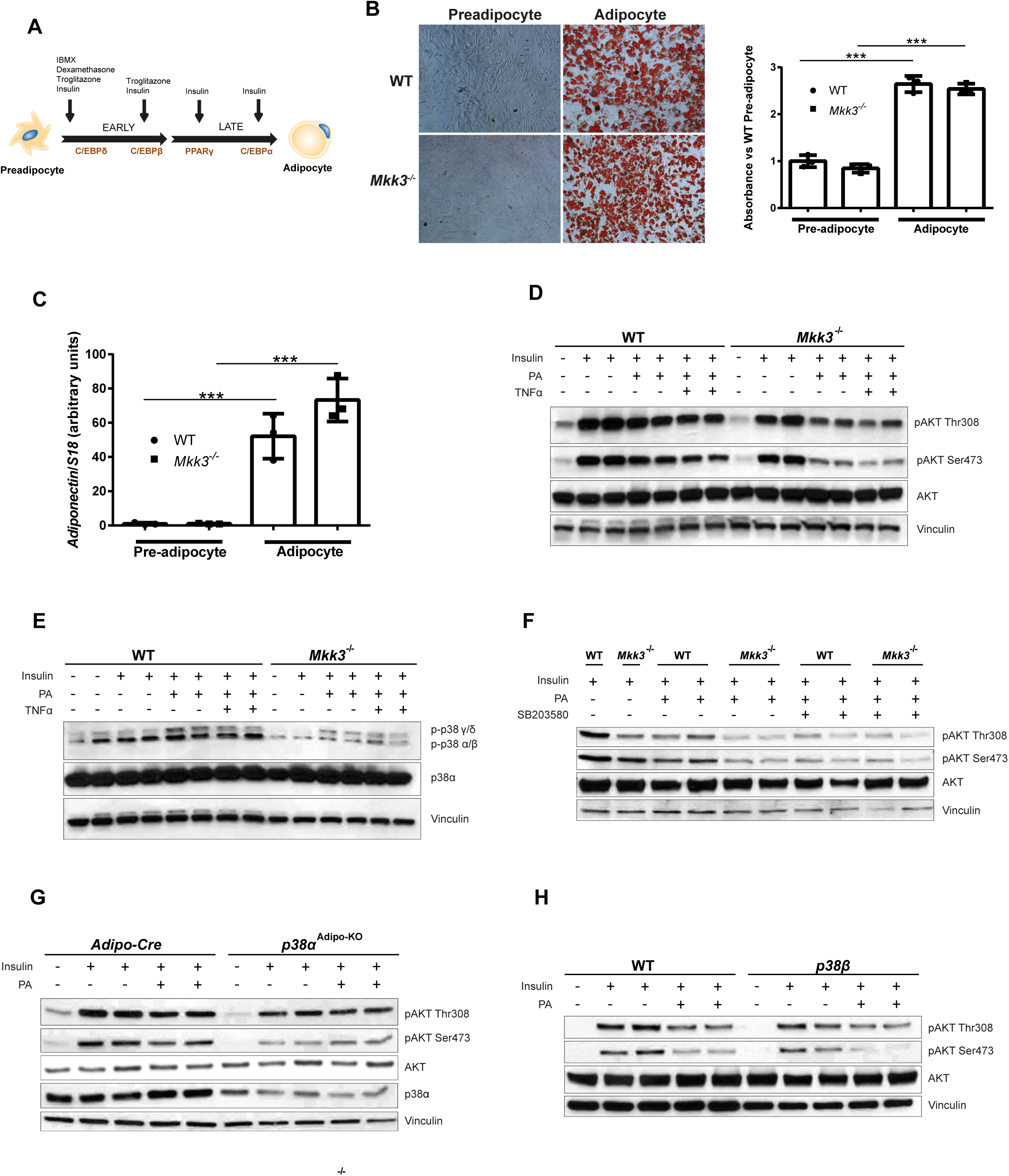
**A**. Scheme representing mature primary adipocyte protocol of differentiation. **B.** Representative images of Oil Red O staining (left) and quantification (right bar plot) of differentiated *Mkk3^-/-^* derived pre-adipocytes showing no differences in lipid accumulation. The data are presented as the colorimetric quantification. **C.** Adipocyte-specific *Adiponectin* expression was not altered in *Mkk3^-/-^*derived mature adipocytes. n= 3 replicates of 3 independent experiments, mean ± SEM, Student’s *t*-test coupled with Bonferroni post-test, ***p<0.001. **D.** Western blot showing reduced activation of AKT in *Mkk3^-/-^* derived adipocytes treated with 0.5mM of palmitate and 10ng/ml TNFα. **E.** Immunoblot showing reduced phosphorylation of p38α/β in *Mkk3^-/-^* derived adipocytes after treatment with 0.5mM of palmitate, 10ng/ml TNFα. **F.** Western blot showing that *Mkk3^-/-^* adipocytes AKT activation is not further reduced when treated for 24h with 10µM p38α/β specific inhibitor SB203580, palmitate 0.5mM. **G.** Immunoblot showing reduced AKT phosphorylation in mature adipocytes derived from white adipose tissue-specific p38α knockout mice after 0.5mM palmitate treatment for 24h. **H.** Western blot showing reduced AKT activation in mature p38β^-/-^ mice derived adipocytes after 24h of 0.5 mM palmitate treatment. D-H, Adipocytes starved for 2h were stimulated or not with 10nM of insulin for 10 min.

To evaluate insulin signaling in *Mkk3^-/-^* differentiated adipocytes, Akt phosphorylation was measured after treatment with obesogenic stimuli of 0.5 mM palmitate (PA) and/or 10ng/ml ng/ml TNFα, followed by stimulation with insulin. p-Akt was induced by insulin stimulation in both WT and *Mkk3^-/-^* adipocytes, although *Mkk3^-/-^* adipocytes displayed lower insulin-induced AKT phosphorylation (Fig. 4D). PA or PA with TNF-α treatment resulted in decreased insulin-stimulated AKT phosphorylation at both residues in WT adipocytes. However, the reduction in AKT phosphorylation was more pronounced in *Mkk3^-/-^* adipocytes, indicating that *Mkk3^-/-^* adipocytes were more insulin-resistant than WT adipocytes (Fig. 4D), which was consistent with the data observed in HFD-fed *Mkk3^-/-^* and *Mkk3*^Fabp4-KO^ adipose tissue, supporting the hypothesis that MKK3 controls insulin sensitivity in an adipocyte-specific manner.

### Insulin resistance in *Mkk3^-/-^* mice depends on the downstream kinases p38α and p38β

It has been recently described that in the heart, MKK3 is the main activator of p38α and p38β, whereas MKK6 mainly regulates p38γ and p38δ (19). Therefore, p38 phosphorylation in adipocytes lacking *Mkk3* upon insulin stimulation was analyzed. Under basal conditions, non-insulin-stimulated *Mkk3^-/-^* adipocytes presented reduced levels of all p38 MAPK isoforms compared to WT (Fig. 4E). Upon insulin stimulation, WT adipocytes showed a slight increase in p38 MAPK activation, especially of p38α and p38β family members (corresponding to the lower band in Fig. 4E), whereas MKK3*-*deficient adipocytes exhibited decreased insulin-stimulated p38 MAPK phosphorylation compared to WT adipocytes.

PA or PA + TNFα treatment in WT adipocytes resulted in increased phosphorylation of all p38, including p38α and p38β (lower band) and p38γ and p38δ (upper bands), whereas MKK3-deficient adipocytes showed a dramatic reduction in phosphorylation of all p38, especially p38α and p38β, when treated with the same stimuli (Fig. 4E). These data suggest that the insulin resistance phenotype observed in *Mkk3^-/-^* adipocytes might be explained by the reduced phosphorylation of p38α and p38β observed in *Mkk3^-/-^* adipocytes.

To address whether the reduced activation of p38α and p38β in *Mkk3*^-/-^ adipocytes was responsible for the insulin resistance phenotype, PA-stimulated WT and *Mkk3^-/-^* adipocytes were treated with the p38α and p38β specific inhibitor SB203580 and insulin-mediated Akt phosphorylation was assessed (Fig. 4F). Inhibition of p38α and p38β by SB203580 together with PA treatment resulted in decreased insulin-induced phosphorylation of Akt in WT adipocytes compared to that in WT adipocytes treated only with PA. However, no further reduction in insulin-stimulated Akt phosphorylation was observed in *Mkk3^-/-^* adipocytes treated with PA or SB203580 compared with that in *Mkk3^-/-^* adipocytes treated only with PA (Fig. 4F). These data suggest that the specific reduction in p38α and p38β activation in *Mkk3^-/-^* adipocytes might be responsible for impaired insulin signaling.

To investigate the role of p38α and p38β in insulin resistance, insulin signaling was evaluated in p38α and p38β knockout adipocytes. In adipocytes lacking p38α, control Adipo-Cre adipocytes showed pronounced insulin-induced AKT phosphorylation, which was strongly reduced upon treatment with PA (Fig. 4G). However, *p38α*^Adipo-KO^ adipocytes displayed reduced insulin-stimulated AKT phosphorylation, which surprisingly was not further reduced by PA treatment. These data might indicate that the lack of p38α is sufficient to produce maximum inhibition of insulin signaling. Although the p38α deletion appeared incomplete owing to the nature of the *Cre* recombinase system used, the level of deletion was sufficient to induce the insulin resistance phenotype (Fig. 4G).

However, when *p38β*^-/-^ primary adipocytes were tested, it was observed that while WT adipocytes presented characteristic insulin-stimulated phosphorylation of AKT, which was reduced upon PA treatment, *p38β*^-/-^ adipocytes displayed reduced basal insulin-stimulated AKT basal phosphorylation compared to WT adipocytes, and insulin-stimulated Akt phosphorylation was further reduced when *p38β*^-/-^ adipocytes were treated with PA (Fig. 4H). This indicates that p38β-deficient adipocytes are more insulin-resistant, and that this insulin resistance is further exacerbated by PA treatment.

These results suggest that, first, insulin resistance observed in *p38α*^Adipo-KO^ and *p38β^-/-^* adipocytes correlated with the phenotype observed in *Mkk3^-/-^* adipocytes; second, ablation of p38α and p38β or reduced phosphorylation of p38α and p38β detected in *Mkk3^-/-^* adipocytes might explain the observed insulin resistance phenotype.

### *p38β* in human visceral fat negatively correlates with body mass index

As the deletion of p38α in adipose tissue results in the protection of insulin resistance (9), we focused on the involvement of p38β. To further investigate the role of the MKK3-p38 axis in obesity and diabetes, *p38β* (*MAPK11)* mRNA was quantified in visceral adipose tissue from a cohort of 71 subjects. We observed that p38β expression was negatively correlated with BMI and circulating C-peptide and leptin levels in plasma (Fig. 5A-C), unlike *MKK3,* which showed positive correlation. Moreover, *MKK3* negatively correlated with *MAPK11* (*p38β*) expression levels in human visceral adipose tissue (Fig. 5D), suggesting that *P38β* expression may be reduced in obesity and that, in fact, *MKK3* overexpression might represent a protective mechanism to compensate for the loss of this kinase.

**Fig. 5.**
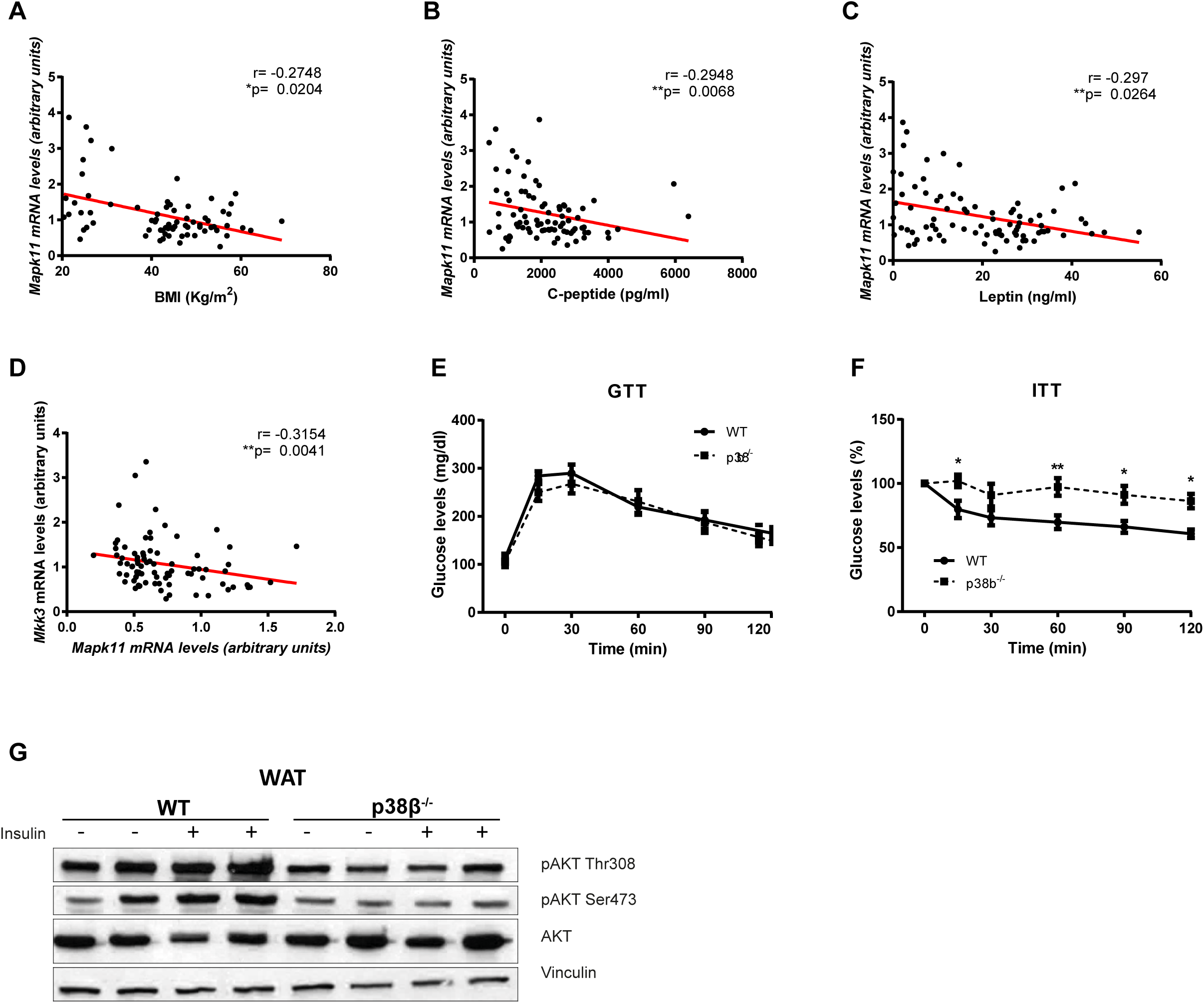
**A-D.** *Mapk11* (*p38β*) expression in visceral adipose tissue from lean and obese human subjects is negatively correlated with BMI (A), circulating levels of C-peptide (B), and leptin (C). **D.** *Mkk3* mRNA expression levels negatively correlates with *Mapk11* in human visceral adipose tissue. Data are presented as discrete value comparisons between individual and regression slope calculated, n= 71, Spearmańs Rho, *p<0.05, **p<0.01. **E.** GTT of HFD-fed WT and *p38β*^-/-^ mice. **F.** ITT of HFD-fed WT and *p38β*^-/-^ mice, ITT indicates that *p38β*^-/-^ mice present increased insulin resistance. **G.** Western blot showing reduced basal and 10nM insulin stimulated for 10 min AKT phosphorylation in white adipose tissue from *p38β*^-/-^ mice.

### HFD-fed p38β deficient mice present increased insulin resistance

To further investigate the role of the MKK3-p38 axis in insulin resistance *in vivo*, mice lacking p38β (*p38β^-/-^*) were fed HFD. In *p38β^-/-^* mice, no differences in glucose tolerance were observed (Fig. 5E), and insulin-stimulated glucose clearance was reduced compared to that in WT mice, indicating higher insulin resistance in *p38β^-/-^* mice (Fig. 5F). Next, insulin signaling in WAT was analyzed, revealing that WAT from *p38β^-/-^* mice showed reduced AKT phosphorylation both under basal conditions and upon insulin stimulation compared to WT mice. This finding indicates that, similar to both germline and adipocyte-specific *Mkk3* KO mice, WAT from HFD-fed *p38β^-/-^*mice was more resistant to insulin than WAT from WT animals (Fig. 5G).

Histological evaluation of WAT showed an increased presence of infiltrated leukocytes and smaller adipocyte size in HFD-fed *p38β^-/-^ mice* compared to WT mice (Fig. S4A), with no differences in the liver and BAT between genotypes. Additionally, to further investigate whether the exacerbated insulin resistance in *p38β^-/-^* mice was due to the absence of *p38β* in leukocytes, chimeric mice were generated by transplanting BM from WT or *p38β^-/-^* donors into lethally irradiated WT recipients, from now referred as WT^WT^ or WT*^p38β-/-^*mice, respectively (Fig. S4B) and fed an HFD. No differences in weight gain or insulin sensitivity were observed between the WT^WT^ and WT*^p38β-/-^ mice* (Fig. S4C, D), suggesting that p38β deficiency in the BM did not affect HFD-induced insulin resistance.

### mTORC1 and S6K activation negatively regulates insulin signaling in *Mkk3^-/-^* adipocytes

Insulin binding to the insulin receptor (IR) in adipose tissue recruits and induces the phosphorylation of Tyr residues of insulin receptor substrate-1 (IRS-1), triggering insulin signaling. In contrast, FFAs, high glucose, and diacylglycerol (DAG) negatively regulate IRS-1 (29, 30), promoting its inactivation by inducing the phosphorylation of Ser/Thr residues by Ser/Thr kinases and blocking Tyr phosphorylation. Therefore, we evaluated whether the lack of MKK3 affects insulin-induced IRS-1 serine phosphorylation in adipocytes. PA-treated *Mkk3^-/-^*adipocytes showed increased phosphorylation of IRS-1 at Ser307 in the basal state (non-insulin-stimulated) compared with controls (Fig. 6A). As S6K, upon upstream activation by mTORC1, phosphorylates IRS-1 at Ser307 and blocks its action, we investigated S6K phosphorylation and activation, as well as its direct substrate ribosomal protein S6. Analysis of PA-treated adipocytes under basal conditions (non-insulin-stimulated) showed S6K phosphorylation and activation, as indicated by increased S6 phosphorylation at Ser240/244 in *Mkk3^-/-^* adipocytes (Fig. 6B), indicating that insulin resistance in *Mkk3^-/-^* adipocytes might be mediated by increased S6K activation and, consequently, higher serine phosphorylation of IRS-1. To evaluate whether insulin resistance in *Mkk3^-/-^*and *p38β^-/-^* adipocytes shares the same molecular mechanism, the phosphorylation and activation of S6K in *p38β^-/-^* adipocytes was studied. Similar to *Mkk3*^-/-^ adipocytes, PA-treated *p38β^-/-^* adipocytes presented increased S6K and S6 phosphorylation compared to WT adipocytes (Fig. 6C), correlating both results.

**Fig. 6.**
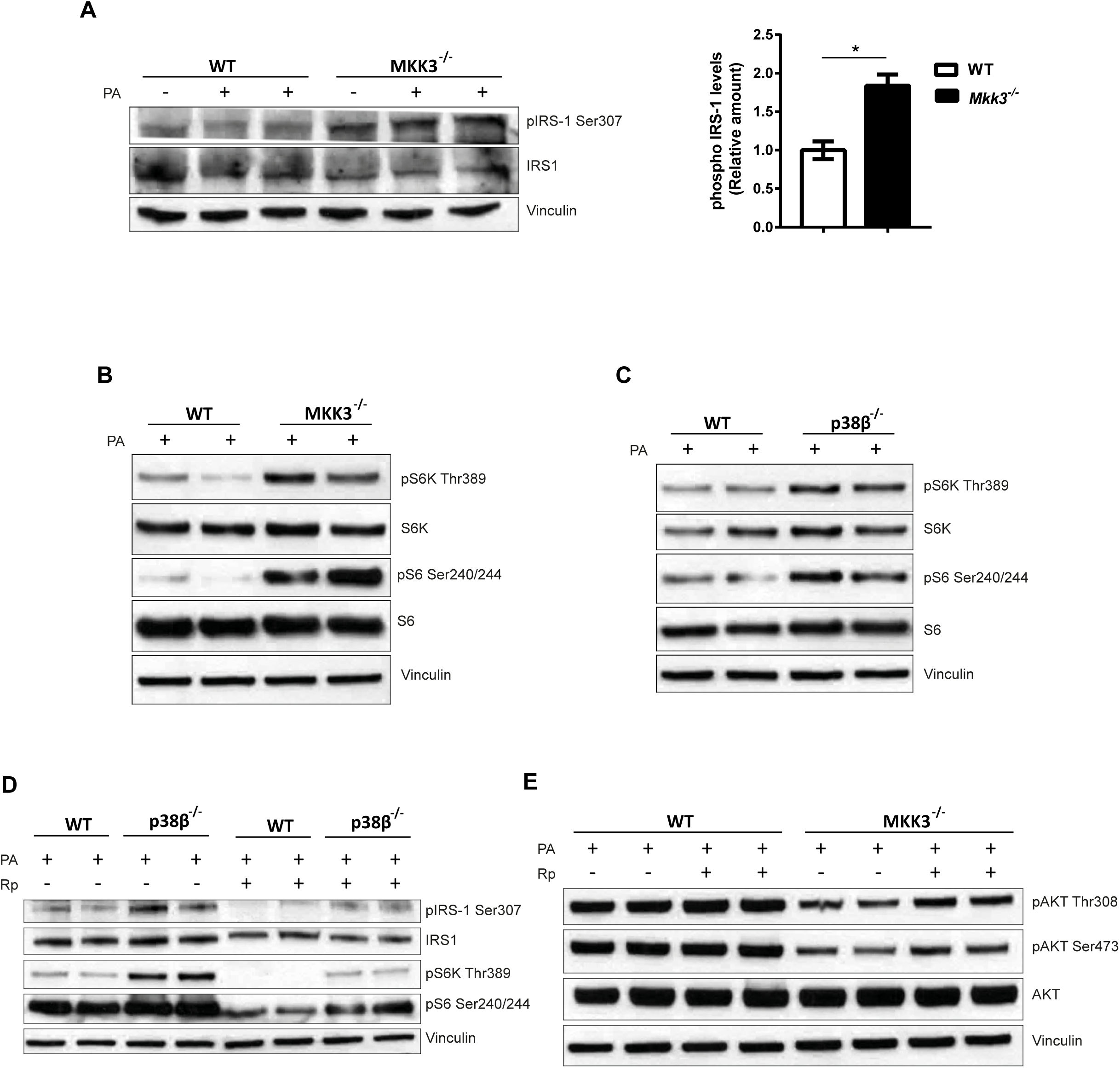
**A.** Immunoprecipitation immunoblot (left) and quantification (right bar plot) showing increased phosphorylation of IRS1 in basal and 0.5mM palmitate stimulated *Mkk3^-/-^* derived adipocytes. Data are presented as mean ± SEM, n=3, Student’s *t*-test coupled to the Bonferroni post-test. *p<0.05. **B, C,** Western blotting showing increased phosphorylation of pS6K and downstream target pS6 in adipocytes derived from *Mkk3^-/-^* (B) and *p38β^-/-^* (C) mice treated with palmitate (0.5mM). **D,** Immunoprecipitation of pIRS-1 Ser307 and western blot showing increased phosphorylation of IRS1, pS6K, and pS6 in *p38β^-/-^* derived adipocytes and partial reversion upon treatment with 10µM mTOR inhibitor rapamycin. **E.** Immunoblot showing partial restoration of AKT phosphorylation in *Mkk3^-/-^* derived adipocytes treated with 10µM of rapamycin.

To assess whether increased S6K activation was involved in the development of insulin resistance, WT and *p38β^-/-^* adipocytes were treated with the mTOR inhibitor rapamycin. PA-treated *p38β^-/-^*adipocytes showed increased phosphorylation of IRS-1 at Ser307 compared to WT adipocytes, correlating with increased S6K activation (Fig. 6D). As expected, rapamycin treatment of WT adipocytes reduced S6K phosphorylation, correlating with the reduced IRS-1 Ser307 phosphorylation, suggesting that S6K is important for PA-induced IRS Ser307 phosphorylation. Interestingly, PA-treated *p38β^-/-^* adipocytes treated with rapamycin showed reduced S6K activation to the same level as WT non-rapamycin-treated WT adipocytes, which correlated with the same level of IRS1 Ser307 phosphorylation in *p38β^-/-^* adipocytes as that in WT adipocytes (Fig. 6D). These results indicate that the MKK3/p38β signaling cascade inhibits S6K phosphorylation in adipocytes, thereby protecting them from obesity-induced insulin resistance.

We previously showed that both PA-treated *Mkk3^-/-^* and *p38β^-/-^* adipocytes showed increased inhibitory phosphorylation of IRS-1 at Ser307, mediated by increased S6K activation; however, the impact of S6K activity on insulin signaling was absent. PA-treated *Mkk3^-/-^* adipocytes showed decreased insulin-stimulated phosphorylation of AKT compared to WT adipocytes (Fig. 6E); however, upon rapamycin treatment, insulin-stimulated Akt phosphorylation in PA-treated Mkk3-/- adipocytes was partially restored, indicating that insulin signaling in *Mkk3^-/-^* adipocytes might be negatively regulated by increased mTORC1 and S6K activation (Fig. 6E).

## Discussion

Here, we investigated the role of the MKK3-p38 axis in the development of HFD-induced insulin resistance. Surprisingly, our results suggest that the activation of MKK3 plays a protective role against insulin resistance in white adipose tissue (WAT). Specifically, we found that HFD-fed Mkk3^-/-^ mice displayed higher insulin resistance than HFD-fed wild-type (WT) mice, owing to increased insulin resistance in WAT. This phenotype was adipocyte-autonomous, as HFD-fed mice lacking MKK3, specifically in adipose tissue (Mkk3^Fabp4-KO^), displayed the same insulin resistance phenotype. In addition, we ruled out the contribution of myeloid cells to the phenotype of Mkk3^Fabp4-KO^ mice, because mice lacking MKK3 in myeloid cells showed no differences in weight distribution or insulin resistance. The absence of p38α, p38γ, and p38δ in the myeloid compartment has been demonstrated to protect against insulin resistance and fatty liver disease (12, 15). In contrast, deletion of MKK3/6 in myeloid cells results in obesity and insulin resistance (18). The lack of a similar phenotype in Mkk3^Lyzs-KO^ mice further suggests that MKK6 may compensate for MKK3 deficiency in myeloid cells but not in adipocytes.

This unexpected finding that HFD-fed Mkk3^-/-^ mice exhibited increased WAT insulin resistance is intriguing because MKK3 activates p38 kinases, which play a crucial role in the development of insulin resistance in various cell types, including hepatocytes and skeletal muscle cells (31, 32). However, recent evidence has demonstrated that MKK3 and MKK6 do not equally activate different members of the p38 family. For example, in muscle cells, MKK3 predominantly activates p38γ and p38δ, whereas MKK6 phosphorylates p38α and p38β (19).

This highlights that p38 activation and insulin resistance may be tissue- and kinase-dependent. This effect of MKK3 deletion on insulin resistance also seems to be independent of the pro-adipogenic effect of p38 activation, as mice lacking MKK3 showed no apparent changes in adipogenesis. Downstream p38 signaling induced by MKK3/6 has been implicated in adipogenesis, but evidence suggests both the induction and inhibition of this process (24). While constitutively active MKK6 induces spontaneous differentiation (33), depletion of p38α promotes adipogenesis through activation of C/EBPβ and peroxisome proliferator-activated receptor-γ (PPARγ) (32). To further analyze the role of the MKK3-p38 axis in insulin sensing by adipocytes, we evaluated signaling in adipocytes lacking p38α and p38β. While adipocytes lacking p38α showed a basal decrease in insulin-induced AKT phosphorylation that was surprisingly not exacerbated by PA treatment, p38β deficiency resulted in a slight decrease in insulin-induced AKT phosphorylation, which was further reduced by treatment with PA. These data suggest that lack of p38α is sufficient to induce maximal inhibition of AKT phosphorylation under basal conditions, whereas p38β improves insulin sensitivity after PA treatment.

Integrating our results mechanistically, we found that MKK3 depletion in adipose tissue leads to reduced activation of p38α and p38β, which in turn induces hyperactivation of S6K. This hyperactivation promotes IRS-1 serine phosphorylation, ultimately leading to insulin resistance (34). Consistent with the importance of mTORC1/S6K activation as a driver of insulin resistance in MKK3 or p38β deficient animals, mTORC1 inhibition by rapamycin was sufficient to partially restore insulin signaling.

Our study results in mouse models align with observations in a human cohort, indicating that upregulation of MKK3 might have a positive impact on obesity and diabetes. This compensatory mechanism may counteract the loss of p38α and p38β in obesity. Our analysis revealed that MKK3 mRNA expression in adipose tissue positively correlated with BMI and circulating C-peptide and leptin levels, whereas P38β (MAPK11) mRNA expression in visceral adipose tissue negatively correlated with BMI and serum C-peptide and leptin levels. Furthermore, MKK3 expression in adipose tissue was negatively correlated with P38β (MAPK11) expression levels in human visceral adipose tissue. As we found that p38β expression levels are reduced in obesity, the increased expression of MKK3 might represent a protective mechanism to compensate for the loss of this kinase.

These human findings, as well as the potential to restore insulin signaling through pharmacological inhibition of the MKK3-p38 pathway, open new avenues for the development of potential therapeutic targets for treating diabetes and obesity. We previously reported that inhibiting p38α might result in protection against diabetes (9); however, this protection may be mediated by the activation of other p38 family members. Therefore, it is essential to clarify the specific role of each family member, as pharmacological inhibitors may exert distinct effects on individual kinases, depending on the target tissue.

## Materials and methods

### Human studies

The study population included obese adult patients with a body mass index (BMI) ≥ 35 recruited from patients who underwent elective bariatric surgery at the Hospital Universitario de Salamanca. Patients were excluded if they had a history of alcohol use disorders, excessive alcohol consumption (>30 g/day in men and > 20 g/day in women), or chronic hepatitis C or B. Control subjects were recruited from patients who underwent laparoscopic cholecystectomy for gallstone disease. This study was approved by the Ethics Committee of the Hospital Universitario de Salamanca, and all participants provided written informed consent to undergo visceral fat biopsy under direct vision during surgery. Data were collected on demographic information (age, sex, and ethnicity), anthropomorphic measurements (BMI), smoking and alcohol history, coexisting medical conditions, and medication use. Before surgery, fasting venous blood samples were collected to measure the complete cell blood count, total bilirubin, aspartate aminotransferase (AST), alanine aminotransferase (ALT), total cholesterol, high-density lipoprotein (HDL), low-density lipoprotein (LDL), triglycerides, alkaline phosphatase (AP), and gamma-glutamyl transpeptidase (GGT) levels.

### Animal models

MKK3 deficient mice (*Mkk3*^-/-^, B6.129-*Map*2*k*3^tm1Flv^) were generated as previously described (35) and kindly donated by Dr. Roger J Davis from the University of Massachusetts. Animals were maintained on a C57BL/6J background (back-crossed 10 generations) and genotype was confirmed by PCR analysis of genomic DNA.

To generate tissue specific knockout mice, we used the *Cre-LoxP* system discovered in the bacteriophage P1 (36, 37). This technique is based in the property of the *Cre* recombinase enzyme of recombining its target *LoxP* sequences, which are placed flanking the gene of interest. *Cre* recombinase recognizes and recombines *the LoxP* sequences and deletes this gene. *Cre* recombinase can be tissue-specific expressed using tissue-specific promoters, such as *Fabp4, Adipoq* (for adipose tissue)*, and Lyz2* (for myeloid cells).

Mice lacking MKK3, specifically in the adipose tissue, were generated as follows. Mice with a germline mutation in Map2k3 and LoxP elements inserted into two introns (Map2k3LoxP) were generated after homologous recombination in ES cells. ES cells were electroporated with a vector and selected using 200 μg/ml G418 and 2 μM ganciclovir. Several correctly targeted ES cell clones were identified by Southern blot and PCR. These ES cell clones were injected into C57BL/6J blastocysts to create chimeric mice that transmitted the mutated Map2k3 allele through the germline. The Flp NeoR cassette was excised by crossing these mice with ACTB:FLPe B6;SJL mice, which express an FLP1 recombinase gene under the direction of the human ACTB promoter. These animals were crossed with mice expressing *Cre* recombinase under the control of the adipose tissue-specific *Fabp4* promoter (*Fabp4-Cre*) (38) on a C57BL/6J background to generate mice lacking MKK3 in adipose tissue. To reduce the number of *LoxP* sequence recombinations needed and achieve better depletion of *the Mkk3* gene, we crossed these mice with full knockout *Mkk3* mice (*Mkk3^-/-^*) to obtain *Fabp4-Cre^+^Mkk3^f/-^* mice (F^KO^ mice). Littermates without the conditional *Mkk3* allele (*Fabp4-Cre^+^Mkk3^+/−^*) were used as controls (F^WT^). Genotype was confirmed by PCR analysis of genomic DNA. Alternatively, mice lacking MKK3 specifically in adipose tissue were generated by crossing *Mkk3^f/f^*mice with mice expressing *Cre* recombinase under the control of the adipose-specific *Adipoq* promoter (*Adipoq-Cre*) (39). To reduce the number of *LoxP* sequence recombinations needed and achieve better depletion of *the Mkk3* gene, these mice were crossed with full knockout *Mkk3* mice (*Mkk3^-/-^*) to obtain *Adipoq-Cre^+^Mkk3^f/-^* mice (Adipo^KO^ mice). Littermates without the conditional *Mkk3* allele (*Adipo-Cre^+^Mkk3^+/−^*) were used as controls (Adipo^WT^). Genotype was confirmed by PCR analysis of genomic DNA. Mice lacking MKK3 specifically in the myeloid lineage tissue were generated by crossing *Mkk3^f/f^*mice with mice expressing *Cre* recombinase under the control of myeloid lineage-specific *Lyz2* promoter (*Lyz2-Cre*), as previously described (40). These mice were also crossed with full knockout *Mkk3* mice (*Mkk3 ^−/−^*) to obtain *Lyz2-Cre^+^Mkk3^f/−^* mice (Lyz^KO^ mice). Littermates without the conditional *Mkk3* allele (*Lyz2-Cre^+^Mkk3^+/−^*) were used as controls (Lyz^WT^). Genotype was confirmed by PCR analysis of genomic DNA. Mice lacking p38β in the whole body were kindly donated by Dr. Angel Nebreda from IRB Barcelona, and were originally generated as p38α^f/f^ p38β^-/-^, and p38α^f/f^ p38β^+/+^ were used as controls.

All mice were housed in a specific pathogen-free (SPF) or quarantine animal facility on a 12-hour light/dark cycle at constant temperature and humidity at the Centro Nacional de Investigaciones Cardiovasculares (CNIC), Universidad de Santiago de Compostela, and Institut de Recerca de Barcelona (IRB), supervised by veterinarian staff. All mice were maintained on a C57BL/6J background (back-crossed 10 generations). All animal experiments conformed to the EU Directive 2010/63EU and Recommendation 2007/526/EC, enforced in Spanish law under Real Decreto 53/2013.

Radiation chimeras using congenic C57BL/6J, *Mkk3*^-/-^, or p38β^-/-^ (CD45.2) and B6. SJL (CD45.1) mice were generated by exposure 8 weeks old B6. SJL recipient mice were administered two doses of ionizing radiation (650 Gy) and reconstituted with 2 × 107 cells from either WT, *Mkk3*^-/-^, or p38β^-/-^ donor BM via tail vein injection. After BM injection, mice were treated with antibiotics in the drinking water *ad libitum*, 1.5 mg/ml gentamicin (Sigma), 1.5 mg/ml kanamycin (Sigma) and 1.5 mg/ml ampicillin (Sigma) for two months. All animals were weighed, fasted for 16 h, bled from the cheek, and stimulated with or without 1.5 U/Kg of insulin injected intraperitoneally for 10 min before sacrifice by cervical dislocation. Tissues were weighed and collected for different purposes, and the tibia was measured using a digital caliper (ratio 6369 HS) relative to organ weight to tibial length. For protein extraction, tissues were snap-frozen in liquid nitrogen (LN2); for RNA isolation, samples were submerged in RNAse inhibitor solution (RNAseZAP, Thermo Fisher Scientific) prior to LN2 snap-freezing; for histological evaluation, tissues were fixed in 10% formalin solution or included in O.C.T (TissueTek) and snap-frozen in LN2.

### Study reconstitution efficiency by flow cytometry

To check the implementation of donor BM and proper depletion of recipient BM in radiation chimera experiments, the stromal vascular fraction (SVF) was isolated by Type-A Collagenase (Roche) disaggregation for 45 min at 37° in an end-to-end water bath, and cellular stromal precipitation was stained using anti 1:200 concentration of CD45.2 and CD45.1 conjugated with PerCP-5.5 and Pacific Blue fluorophores (BD Biosciences), respectively, and analyzed by flow cytometry.

### Obesity-induced diabetic model

Diabetes is an obesity-related disease; hence, feeding mice a fat-enriched diet is an established model to study metabolic disorders that resemble metabolic syndrome in humans (41).

Adult mice (8–12 weeks old) *were fed ad libitum* with a fat-enriched diet (HFD), in which 60% of the calories were fat-derived and 1.5% cholesterol (Research Diets) for 8–12 weeks. The mice were weighed every two weeks to evaluate weight gain.

### In-vivo metabolic tests

Several metabolic parameters were analyzed in the Animal Facility at the CNIC or the Animal Facility at the Universidad de Santiago de Compostela. The Glucose Tolerance Test (GTT) was performed as described previously (5) and allowed us to assess glucose metabolism status in animals. Mice were starved for 16h and basal glucose levels were measured by collecting one blood drop from the distal part of the tail and using a glucometer and glucose strips (Breeze2, Bayer). After basal measurement, we injected i. p. 1 mg/g of glucose (Sigma) dissolved in 1X PBS and detected glucose at 15, 30, 60, 90, and 120 min post-injection. Glucose levels were expressed in mg/dl. The insulin tolerance test (ITT) was performed as previously described (5), which allowed us to determine how peripheral tissues respond to insulin and whether these tissues are sensitive or resistant to insulin. ITT is mechanically performed similarly to GTT, however animals were starved just for 1h, when basal glucose level was quantified, and then 0.75 U/Kg of insulin injected. We used the same time points as the GTT: 15, 30, 60, 90, and 120 min after insulin injection. Glucose levels were relative to the basal glucose levels and expressed as percentages (%).

Magnetic Resonance Imaging (MRI) was performed by Dr. Rubén Nogueiras Laboratory at the Universidad de Santiago de Compostela. WT and *Mkk3*^-/-^ mice were fed an HFD for 8 weeks and transferred to Santiago de Compostela, where they were acclimatized for 1 week.

After acclimatization, fat mass and lean mass were measured by MRI (Whole Body Composition Analyzer; EchoMRI, Houston, TX, USA). Fat mass and lean mass values were expressed as mass (g) and percentage (%), respectively, relative to the full body weight.

### Cell culture

In this study, we isolated and cultured pre-adipocytes and differentiated them into mature adipocytes.

Subcutaneous fat depots were surgically collected and submerged in Dulbeccós Modified Eagle Medium (DMEM, Gibco). Adipose tissue was first mechanically disaggregated using tweezers and razors and then enzymatically disaggregated using Type-A Collagenase medium containing 2 mg/ml collagenase type-A (Roche), 20 mg/ml BSA (Sigma) and 1X PBS (2 ml of solution per animal). Fat tissue was incubated with collagenase solution at 37 °C in a rotating water bath. After disaggregation, the reaction was stopped by adding three volumes of DMEM supplemented with F12 (Gibco), 10% Fetal Bovine Serum (FBS) (Lonza), 2mM L-Glutamine (Sigma), and 100 U/ml penicillin/streptomycin (Sigma). The cell suspension was passed through a 70µm cell strainer (Falcon) to eliminate stroma and debris and centrifuged at 250G for 8 min at RT. The pellet was collected and stored, and the supernatant was re-centrifuged at 400G for 8 min at RT. Both pellets were combined, and cells were counted using a CasyTon cell counter.

Equal numbers of cells were seeded on 10 cm cell culture plates (Costar Corning) and placed in an incubator at 37 °C with 5% CO_2_.

For pre-adipocyte differentiation, when the pre-adipocytes reached confluence, differentiation was initiated by adding DMEM-F12 medium supplemented with 10% FBS, L-glutamine, penicillin/streptomycin, and a cocktail of differentiating compounds composed of 5 µg/ml bovine insulin (Sigma), 1µg/ml dexamethasone (Sigma), 25µg/ml µg/mL 3-isobutyl-1-methylxanthine (IBMX) (Sigma), and 1µM troglitazone (Tocris). On the first day of differentiation, all the indicated compounds were added for 48 h. The medium was then replaced with fresh medium containing insulin and troglitazone and incubated for another 48 h. After this, the medium was changed again and insulin was added. This final step was repeated once more, for a total of 96 hours. After adipocyte differentiation, the cells were starved for 22h in DMEM supplemented with 5% FBS and fully starved for 2h in DMEM without FBS. During 5% FBS starvation and 2h of full starvation, adipocytes were treated with different compounds, depending on the experiment. As obesogenic treatments, 0.5 mM palmitate (Sigma) and 10 ng/ml TNF-α (Sigma) for 24h were used, while 10µM µM SB203580 (ShelleckChem) and 10µM µM rapamycin (ShelleckChem) for 24 h were used to inhibit p38α/β and mTOR, respectively. After treatment, adipocytes were stimulated with 10 nM bovine insulin (Sigma) for 10 min. Untreated insulin-stimulated adipocytes were used as controls. After differentiation, treatment, and stimulation, cells were washed with 1X PBS, extracted according to the posterior analysis, and stored at −80 °C.

### Histology

Liver, WAT, and BAT tissue samples were previously fixed in 10% formalin dilution for at least 48 h, dehydrated in different concentrations of ethanol by the Histology Unit at CNIC, and embedded in paraffin using a Tissue Embedding Center (Leica, EG1150). Paraffin blocks with embedded tissues were cut between 5 and 8 µm thickness using a microtome and dried for 24h at 37°C. Paraffin was removed, and the tissue was hydrated again following sequential 2 min baths of xylene, 100% ethanol, 70% ethanol, 50% ethanol, and water. For hematoxylin/eosin staining, hematoxylin stains acidic structures, such as cellular nuclei, and eosin stains basic structures present in the cytoplasm (42). Slides were hydrated, stained, dehydrated, and mounted in Permount medium (Thermo Fisher Scientific). Slides were observed using an optical microscope (Leica DM2500) at different magnifications, and images were taken using Leica Application Suite (L.A.S).

### Immunofluorescence staining

WAT slides that were previously rehydrated were boiled for 5 min using a pressure cooker and antigen retrieval buffer (10 mM Sodium citrate pH6, 0.1% Tween 20). Slides were permeabilized and endogenous peroxidase was blocked using a solution of 3% methanol in hydrogen peroxide for 30 min at RT. Slides were blocked with 2% Bovine Serum Albumin (BSA) for 1h, washed with 1X PBS, and incubated overnight with anti-F4/80, anti-Ly6G, and anti-perilipin conjugated with Alexa Fluor 647, allophycocyanin (APC), and phycoerythrin (PE) (BD Biosciences), respectively. DAPI was used for nuclear staining for 1 h. Samples were observed, and images were taken using a confocal laser scanning microscope (Leica, TCS-SP5).

### Oil-Red O staining

Oil Red O (Sigma) is a lysochrome that stains neutral triglycerides and lipids in the tissues (43). Oil Red O (0.7%) was dissolved in propylene glycol, filtered, and stored at 55 °C. We used this stain in adipocyte cultures that were previously fixed with 4% paraformaldehyde (16% PFA, Sigma) for 2 h. Adipocytes were dehydrated following sequential baths of propylene glycol and stained with Oil Red O at RT for 2h shaking on a plate shaker. Excess Oil Red O was removed using water and the cells were examined under an optical microscope (Leica DM2500). Images were captured using the Leica Application Suite (L. A. S.).

For quantification, equal volumes of methanol were added to each well to elute the Oil Red O dye, which retains its red color in the solution. Color absorbance was measured using a spectrophotometer (BioRad Benchmark Plus) at 520 nm wavelength and relativized to non-differentiated WT adipocyte absorbance. The adipocyte plates were stored in water at 4 °C for long-term storage.

### Western blot and immunoprecipitation

Total proteins from white adipose tissue (WAT), liver and muscle were extracted in lysis buffer (50 mM Tris-HCl pH 7.5, 1 mM EGTA, 1 mM EDTA pH 8.0, 50 mM Sodium

Fluoride, 1 mM sodium glycerophosphate, 5 mM pyrophosphate, 0.27 M sucrose, 1% Triton X-100, 0.1 mM PMSF, 0.1% β-mercaptoethanol, 1 mM sodium-ortovanadate, 1 µg/ml leupeptin, 1 µg/ml aprotinin). The tissues were homogenized with a Turrax (IKA T-100), centrifuged at 14.000 rpm at 4 °C for 20 min, and the supernatant was collected.

The protein concentration in the supernatants was quantified using the Bradford method This technique is based on a colorimetric reaction: the Coomassie blue reagent (Coomassie G-250, BioRad) turns blue when interacting with proteins containing acidic and aromatic amino acids. This color shift can be measured using a spectrophotometer reading at 595 nm wavelength and extrapolated to a standard curve made with known concentrations of protein solution. Two microliters of our protein solution were added to a 96-well plate (Corning Costar) and 200 µl of Coomassie reagent was used to develop color. The reaction was kept at room temperature (RT) for 5 min, and absorbance was detected using a spectrophotometer (BioRad Benchmark Plus).

Lysates were denatured in loading buffer (250mM Tris-HCl pH 6.8, 10% SDS, 30% glycerol, 0.01% bromophenol blue, and 5% β-mercaptoethanol) at 95 °C for 5 min. Equal amounts of lysates were loaded onto 10% to 12% polyacrylamide gels (30% Acrylamide/Bis Solution, 29:1, Bio-Rad) and run at 150v for 90 min. Gels were transferred to 0.2µm pore nitrocellulose membranes (Bio-Rad), blocked with 10% fat-free milk powder for 45 min, and probed overnight (ON) with the following primary antibodies at 1:1000 concentration: phospho AKT Thr308 (#9275), phospho AKT Ser473 (#9271), AKT (#9272), phospho p38 Thr180/Tyr182 (#9211), phospho S6K Thr389 (#9206), S6K (#2708), phospho S6 Ser240/244 (#5364), S6 (#2217), all from Cell Signaling Technologies, Vinculin (V4505), and p38α (SC-535) from Sigma and Santa Cruz, respectively. Anti-rabbit or anti-mouse antibodies conjugated to horseradish peroxidase secondary antibodies were used at a 1:5000 concentration in BSA 5% and incubated for 1 h at room temperature (RT). Western blots were developed using an ECL detection reagent (Amersham ECL, GE Healthcare).

To improve specificity and signal strength, some proteins were immunoprecipitated using G-Sepharose beads. 20 µl of G-Sepharose beads were incubated with 2µg of IRS-1 antibody, produced in the laboratory. Beads and antibodies were incubated in a laboratory wheel at 4°C for 1h. After incubation, the samples were centrifuged twice with 1X PBS at 14000rpm at RT on a benchtop centrifuge to wash the samples, and one last time with lysis buffer. After washing the beads, 750 µg of protein were added and incubated overnight at 4 °C on a rotating wheel. The next day, the samples were centrifuged, and the unbound fraction was collected and stored at −80 °C. Precipitated beads were washed 3X with lysis buffer and denatured at 95 °C for 5 min in loading buffer. We centrifuged the samples, loaded the supernatant on pre-cast gels 4%-10% acrylamide (Bio-Rad), and transferred it as described in SDS-PAGE electrophoresis. Phospho-IRS-1 Ser307 (07-247) and IRS-1 (#2382) antibodies from Up-state and Cell Signaling Technologies, respectively, were used. **RNA isolation and qPCR.**

mRNA was extracted from tissues and cells following the manufacturer’s instructions (Qiagen RNA Mini Extraction Kit). QIAzol lysis reagent and Turrax (IKA T-100) were used on tissues and cells to homogenize the samples. The mRNA was quantified using a NanoDrop spectrophotometer (ND-1000, Thermo Scientific). For retrotranscription, 100 µg of mRNA either from WAT or adipocytes was transcribed to cDNA using a High-Capacity cDNA Reverse Transcription Kit (Thermo Fisher) following the manufacturer’s instructions. The reaction was performed for 10 min at 25°C, 2h at 337 °Cand 5 min at 85 °C. qPCR was performed using specific primers and the Fast SYBR Green system (Applied Biosystems) or TaqMan Assays (Applied Biosystems) on a 7900HT Fast Real-time PCR Thermal Cycler (Applied Biosystems). A dissociation curve cycle was used after each reaction to verify the primer specificity and purity. Expression levels were normalized to S18 and Actin and mRNA expression levels were calculated by interpolating Ct values on a standard curve of mRNA.

### Luminex circulating cytokine analysis

We analyzed the levels of insulin, C-peptide, adiponectin, and leptin, two circulating cytokines, and two adipokines in serum extracted from blood obtained from lean and obese patients.

Briefly, Luminex is based on microspheres coated with specific antibodies, which are internally dyed with a combination of red and infrared dyes. Different combinations of red and infrared dyes allowed for the specific detection of each microsphere, allowing the detection of multiple analytes in the same sample.

Serum cytokine and adipokine concentrations were measured by multiplex enzyme-linked immunosorbent assay (ELISA) using a Luminex 200 IS analyzer (Millipore), following the manufactureŕs instructions.

### Statistical analysis

Data are presented as mean ± Standard Error of Mean (SEM), and GraphPad 7 and SPSS v20 were used for statistical analysis.

When two groups were examined for statistical significance, we used a Two-tailed Student’s *t-*test. When more than two groups were examined for statistical significance, analysis of variance (ANOVA) coupled with Bonferronís post-test was used. Spearmańs Rho or Pearsońs correlation coefficients were used in human correlation graphs. Statistical differences were considered when the p value was lower than 0.05 (* p<0.05, ** p<0.01, *** p<0.001).

## Conflict of interest

The authors declare that they have no conflicts of interest.

## Acknowledgements

EB is a fellow of the Ministerio de Economía y Competitividad (SAF2010-19347). L. Herrera-Melle is a fellow of the Ministerio de Educación, Cultura y Deporte (FPU15-05802). GS is supported by the following projects: PMP21/00057 Infraestructura de Medicina de Precisión asociada a la Ciencia y Tecnología IMPACT-2021 funded by the Instituto de Salud Carlos III (ISCIII), PDC2021-121147-I00, PID2019-104399RB-I00 funded by MCIN/AEI/10.13039/501100011033, all of them funded by the European Union (FEDER/FSE), “Una manera de hacer Europa”/ “El FSE invierte en tu futuro”/ Next Generation EU and cofunded by the European Union / Plan de Recuperación, Transformación y Resiliencia (PRTR) and PID2022-138525OB-I00 (Agencia Estatal de Investigación 10.13039/501100011033) funded by MICIU/AEI/10.13039/501100011033 fondos FEDER and EU. The following grants provided additional funding: Instituto de Salud Carlos III (ISCIII), PDC2021-121147-I00 Convocatoria: Proyectos Prueba de Concepto 2021 Ministerio de Ciencia e Innovación and PID2022-138525OB-I00 Ministerio de Ciencia e Innovación to GS, Fundación Jesús Serra, EFSD/Lilly European Diabetes Research Programme, BBVA Foundation Leonardo Grants program for Researchers and Cultural Creators (Investigadores-BBVA-2017), Fundación AECC PROYE19047SABI and Comunidad de Madrid IMMUNOTHERCAN B2017/BMD-3733 to GS; PMP21/00113 to JLT and PI20/00743 and INT21/00065 to MM (funded by ISCIII and cofunded by the European Union), Junta de Castilla y Leon GRS 2868/A2/2023 to JLT and GRS 2388/A/21, GRS 2648/A/22, and GRS 2916/A1/2023 to MM; Sociedad Española de Medicina Interna (SEMI 2024) and Sociedad Castellano-Leonesa Cántabra de Medicina Interna (SOCALMI 2022) to MM. GS is a Miembro Numerario of the RACVE. The CNIO is supported by the Instituto de Salud Carlos III (ISCIII), the Ministerio de Ciencia, Innovación y Universidades (MCNU), and is a Severo Ochoa Center of Excellence (SEV-2015-0510).

---

**Supplementary Fig. 1.**

**A**. *Mkk3^-/-^* mice fed an HFD for 12 weeks present reduced fat mass measured by MRI. **B, C.** Decreased white (B) and brown (C) adipose tissue weights relative to tibia length of HFD-fed *Mkk3^-/-^*mice. **D.** Increased lean mass measured using MRI. **E, F.** Unchanged liver weight (E) and increased muscle weight (F) in HFD-fed *Mkk3^-/-^* mice. Data are presented as mean ± SEM, n=10, Student’s *t*-test coupled with Bonferroni post-test, *p<0.05, **p<0.01, ***p<0.001. **G.** Representative H&E images showing increased leukocyte infiltration in white adipose tissue (middle lane images) and normal liver (top lane) and brown adipose tissue (bottom lane) morphology of HFD-fed *Mkk3^-/-^* mice. Scale bar represents 100µm. **H.** Immunofluorescence images of white adipose tissue sections indicated an increased number of F4/80^+^ (red) and Ly6G^+^ (grey) cells and reduced perilipin levels (green). Scale bar represents 50µm. **I-N.** Expression of myeloid markers *Lyz2* (I)*, Emr1* (J), chemoattractants *Ccl2* (K) *and Ccl3* (L), and pro-inflammatory cytokines *Il6* (M) and *Tnf*α (N) remained unchanged in white adipose tissue from HFD-fed *Mkk3^-/-^*mice. Data are represented as mean ± SEM, n= 6-10, Student’s *t*-test coupled with the Bonferroni post-test.

**Supplementary Fig. 2.**

**A**. Schematic of the experimental approach (left) and cytometry analysis (right plot) of acceptor stromal vascular fraction, plots showing more than 95% efficiency of WT or *Mkk3^-/-^*bone marrow reconstitution transplanted into 8 month old lethally irradiated WT recipient congenic mice. **B.** Weight gain after 6 weeks of HFD of WT*^WT^* and WT*^Mkk3-/-^* mice two months after bone marrow transplantation. Data are presented as mean ± SEM (n = 9), Two-way ANOVA coupled with Bonferroni post-test. **C-F.** Unaltered liver (C), muscle (D), white (E), and brown (F) adipose tissue weight differences relative to tibial length in WT*^Mkk3-/-^* mice after six weeks of HFD feeding. Data are presented as mean ± SEM, n=9, Student’s *t-*test coupled with the Bonferroni post-test. **G, H.** GTT and ITT of WT*^Mkk3-/-^* mice fed an HFD for 6 weeks showed no differences in glucose or insulin tolerance. Data are presented as direct measure of glucose in GTT and decrease vs. basal glycemia expressed as percentage in ITT, mean ± SEM, n=9, Two-way ANOVA coupled to Bonferroni post-test. **I.** Unchanged weight gain of myeloid-specific *Mkk3* conditional mice (*Mkk3*^Lyz2-KO^) fed an HFD for 10 weeks. Data are presented as the mean ± SEM, n=9-12, Two-way ANOVA coupled with Bonferroni post-test. **J-M.** No differences in liver (J), muscle (K), white (L), and brown (M) adipose tissue weights were observed in *Mkk3*^Lyz2-KO^ mice. **N.** The insulin tolerance test showed no differences in the insulin sensitivity of *Mkk3*^Lyz2-KO^ mice fed an HFD for 12 weeks. Data are presented as the mean ± SEM, n=9-12, Two-way ANOVA coupled with Bonferroni post-test.

**Supplementary Fig. 3.**

**A-D**. Unchanged weight relative to tibial length of the liver (A), muscle (B), white (C), or brown (D) adipose tissue of HFD-fed *Mkk3*^Fabp4-KO^ mice for 10 weeks. Data are presented as mean ± SEM, n=8-11, Student’s *t*-test coupled with the Bonferroni post-test.

**Supplementary Fig. 4.**

**A**. Representative H&E staining images of the liver, white adipose tissue, or brown adipose tissue of HFD-fed *p38β*^-/-^ mice showing increased leukocyte infiltration in white adipose tissue (middle lane) of *p38β*^-/-^ mice. Scale bar represents 100µm. **B.** Schematic of experimental approach (top cartoon) and (lower plots) Bone marrow transplantation of WT and p38β^-/-^ derived donor mice into lethally irradiated congenic mice show above 94% of efficiency of donor bone marrow reconstitution in stromal vascular fraction analysis from acceptor white adipose tissue. **C.** After 2 weeks, reconstituted mice were fed an HFD for 8 weeks with no differences in weight gain. **D.** ITT show no differences in insulin tolerance. In (C) and (D), data are presented as mean ± SEM, n=8-9, Two-way ANOVA coupled with Bonferroni post-test.

## References

1. R. Divella, R. De Luca, I. Abbate, E. Naglieri, A. Daniele, Obesity and cancer: the role of adipose tissue and adipo-cytokines-induced chronic inflammation. Journal of Cancer 7, 2346–2359 (2016).

2. B. K. Surmi, A. H. Hasty, Macrophage infiltration into adipose tissue: initiation, propagation and remodelling. Future lipidology 3, 545–556 (2008).

3. B. B. Kahn, J. S. Flier, Obesity and insulin resistance. J Clin Invest 106, 473–481 (2000).

4. G. van Hall et al., Interleukin-6 stimulates lipolysis and fat oxidation in humans. J Clin Endocrinol Metab 88, 3005–3010 (2003).

5. G. Sabio et al., A stress signaling pathway in adipose tissue regulates hepatic insulin resistance. Science 322, 1539–1543 (2008).

6. E. Manieri, G. Sabio, Stress kinases in the modulation of metabolism and energy balance. J Mol Endocrinol 55, R11–22 (2015).

7. C. J. Carlson, C. M. Rondinone, Pharmacological inhibition of p38 MAP kinase results in improved glucose uptake in insulin-resistant 3T3-L1 adipocytes. Metabolism: clinical and experimental 54, 895–901 (2005).

8. I. Nikolic et al., Lack of p38 activation in T cells increases IL-35 and protects against obesity by promoting thermogenesis. EMBO Rep 10.1038/s44319-024-00149-y (2024).

9. N. Matesanz et al., p38α blocks brown adipose tissue thermogenesis through p38δ inhibition. PLoS biology 16, e2004455 (2018).

10. C. de Alvaro, T. Teruel, R. Hernandez, M. Lorenzo, Tumor necrosis factor alpha produces insulin resistance in skeletal muscle by activation of inhibitor kappaB kinase in a p38 MAPK-dependent manner. J Biol Chem 279, 17070–17078 (2004).

11. G. Li, E. J. Barrett, M. O. Barrett, W. Cao, Z. Liu, Tumor necrosis factor-alpha induces insulin resistance in endothelial cells via a p38 mitogen-activated protein kinase-dependent pathway. Endocrinology 148, 3356–3363 (2007).

12. X. Zhang et al., Macrophage p38alpha promotes nutritional steatohepatitis through M1 polarization. J Hepatol 71, 163–174 (2019).

13. L. Herrera-Melle et al., p38alpha kinase governs muscle strength through PGC1alpha in mice. Acta Physiol (Oxf) 240, e14234 (2024).

14. C. Folgueira et al., Remodelling p38 signaling in muscle controls locomotor activity via IL-15. Sci Adv 10, eadn5993 (2024).

15. B. Gonzalez-Teran et al., p38gamma and p38delta reprogram liver metabolism by modulating neutrophil infiltration. EMBO J 35, 536–552 (2016).

16. M. Crespo et al., Myeloid p38 activation maintains macrophage-liver crosstalk and BAT thermogenesis through IL-12-FGF21 axis. Hepatology 10.1002/hep.32581 (2022).

17. M. Crespo et al., Neutrophil infiltration regulates clock-gene expression to organize daily hepatic metabolism. Elife 9 (2020).

18. M. Crespo et al., Myeloid p38 activation maintains macrophage-liver crosstalk and BAT thermogenesis through IL-12-FGF21 axis. Hepatology 77, 874–887 (2023).

19. R. Romero-Becerra et al., MKK6 deficiency promotes cardiac dysfunction through MKK3-p38gamma/delta-mTOR hyperactivation. Elife 11 (2022).

20. G. Remy et al., Differential activation of p38MAPK isoforms by MKK6 and MKK3. Cell Signal 22, 660–667 (2010).

21. N. Matesanz et al., MKK6 controls T3-mediated browning of white adipose tissue. Nat Commun 8, 856 (2017).

22. 22. R. Romero-Becerra, et al., p38gamma/delta activation alters cardiac electrical activity and predisposes to ventricular arrhythmia. Nat Cardiovasc Res 2, 1204–1220 (2023).

23. S. C. Ruffolo et al., Basal activation of p70S6K results in adipose-specific insulin resistance in protein-tyrosine phosphatase 1B -/- mice. The Journal of biological chemistry 282, 30423–30433 (2007).

24. I. Nikolic, M. Leiva, G. Sabio, The role of stress kinases in metabolic disease. Nat Rev Endocrinol 16, 697–716 (2020).

25. M. P. Czech, Insulin action and resistance in obesity and type 2 diabetes. Nature medicine 23, 804–814 (2017).

26. W. Liu et al., Serum leptin, resistin, and adiponectin levels in obese and non-obese patients with newly diagnosed type 2 diabetes mellitus: A population-based study. Medicine 99, e19052 (2020).

27. S. Cinti et al., Adipocyte death defines macrophage localization and function in adipose tissue of obese mice and humans. Journal of lipid research 46, 2347–2355 (2005).

28. L. Boutens et al., Unique metabolic activation of adipose tissue macrophages in obesity promotes inflammatory responses. Diabetologia 61, 942–953 (2018).

29. K. D. Copps, M. F. White, Regulation of insulin sensitivity by serine/threonine phosphorylation of insulin receptor substrate proteins IRS1 and IRS2. Diabetologia 55, 2565–2582 (2012).

30. S. Boura-Halfon, Y. Zick, Phosphorylation of IRS proteins, insulin action, and insulin resistance. Am J Physiol Endocrinol Metab 296, E581–591 (2009).

31. Y. Wu et al., Activation of AMPKalpha2 in adipocytes is essential for nicotine-induced insulin resistance in vivo. Nat Med 21, 373–382 (2015).

32. M. Leiva, N. Matesanz, M. Pulgarin-Alfaro, I. Nikolic, G. Sabio, Uncovering the Role of p38 Family Members in Adipose Tissue Physiology. Front Endocrinol (Lausanne) 11, 572089 (2020).

33. J. A. Engelman et al., Constitutively active mitogen-activated protein kinase kinase 6 (MKK6) or salicylate induces spontaneous 3T3-L1 adipogenesis. J Biol Chem 274, 35630–35638 (1999).

34. V. Aguirre et al., Phosphorylation of Ser307 in insulin receptor substrate-1 blocks interactions with the insulin receptor and inhibits insulin action. J Biol Chem 277, 1531–1537 (2002).

35. H. T. Lu et al., Defective IL-12 production in mitogen-activated protein (MAP) kinase kinase 3 (Mkk3)-deficient mice. Embo J 18, 1845–1857 (1999).

36. B. Sauer, N. Henderson, Site-specific DNA recombination in mammalian cells by the Cre recombinase of bacteriophage P1. Proceedings of the National Academy of Sciences of the United States of America 85, 5166–5170 (1988).

37. P. C. Orban, D. Chui, J. D. Marth, Tissue- and site-specific DNA recombination in transgenic mice. Proc Natl Acad Sci U S A 89, 6861–6865 (1992).

38. W. He et al., Adipose-specific peroxisome proliferator-activated receptor gamma knockout causes insulin resistance in fat and liver but not in muscle. Proc Natl Acad Sci U S A 100, 15712–15717 (2003).

39. J. Eguchi et al., Transcriptional control of adipose lipid handling by IRF4. Cell metabolism 13, 249–259 (2011).

40. B. E. Clausen, C. Burkhardt, W. Reith, R. Renkawitz, I. Forster, Conditional gene targeting in macrophages and granulocytes using LysMcre mice. Transgenic Res 8, 265–277 (1999).

41. R. Buettner, J. Scholmerich, L. C. Bollheimer, High-fat diets: modeling the metabolic disorders of human obesity in rodents. *Obesity (Silver Spring*, Md.) 15, 798–808 (2007).

42. A. H. Fischer, K. A. Jacobson, J. Rose, R. Zeller, Hematoxylin and eosin staining of tissue and cell sections. CSH protocols 2008, pdb.prot4986 (2008).

43. G. J. Hausman, Techniques for studying adipocytes. Stain technology 56, 149–154 (1981).

44. M. M. Bradford, A rapid and sensitive method for the quantitation of microgram quantities of protein utilizing the principle of protein-dye binding. Analytical biochemistry 72, 248–254 (1976).

